# Part-night exposure to artificial light at night has more detrimental effects on aphid colonies than fully lit nights

**DOI:** 10.1101/2022.10.24.513514

**Authors:** Robin Heinen, Oriana Sanchez-Mahecha, T. Martijn Bezemer, Davide M. Dominoni, Claudia Knappe, Johannes Kollmann, Anton Kopatsch, Zoë A. Pfeiffer, Michael Schloter, Sarah Sturm, Jörg-Peter Schnitzler, A. Corina Vlot, Wolfgang W. Weisser

## Abstract

Artificial light at night (ALAN) threatens natural ecosystems globally. While ALAN research is increasing, little is known about how ALAN affects plants and interactions with other organisms. We explored the effects of ALAN on plant defence and plant–insect interactions using barley (*Hordeum vulgare*) and the English grain aphid (*Sitobion avenae*). Plants were exposed to ‘full’ or ‘part’ nights of 15–20 lux ALAN, or no ALAN ‘control’ nights, to test the effects of ALAN on plant growth and defence. Although plant growth was only minimally affected by ALAN, aphid colony growth and aphid maturation were reduced significantly by ALAN treatments. Importantly, we found strong differences between full-night and part-night ALAN treatments. Contrary to our expectations, part ALAN had stronger negative effects on aphid colony growth than full ALAN. Defence-associated gene expression was affected in some cases by ALAN, but also positively correlated with aphid colony size, suggesting that the effects of ALAN on plant defences are indirect, and regulated via direct disruption of aphid colonies, rather than via ALAN-induced upregulation of defences. Mitigating ecological side effects of ALAN is a complex problem, as reducing exposure to ALAN increased its negative impact on insect herbivores.

## Introduction

Artificial light at night (ALAN) is increasingly common in urban and rural areas of the world (Gaston, Visser & Hölker, 2015; Kyba et al., 2017). It is emitted by various man-made sources including street lights, building lights, and advertising lights. Current levels of ALAN are unprecedented in the ecological history of life on Earth, as light levels followed a predictable periodicity of diurnal and annual rhythms, with little light at night except of stars and the moon. These rhythms are increasingly disrupted by ALAN in many areas of the world, and ALAN therefore is considered an important aspect of global change (Gaston, Visser & Hölker, 2015; Senzaki et al., 2020; Falcón et al., 2020). For instance, ALAN can interfere with natural behaviour and physiology in many taxa (Sanders et al., 2021), including freshwater and marine invertebrates (Duarte et al. 2019; Perkin et al. 2011; Ayalon et al. 2019), terrestrial insects and several vertebrates, such as bats and birds (Desouhant et al., 2019; Stone, Harris & Jones, 2015; Dominoni, Quetting & Partecke, 2013). Surprisingly, given their dependence on the light environment for acquisition of energy, and as signal for photomorphogenesis, phenological and defence processes, much less is known about how ALAN affects plants (Briggs 2006; Bennie et al. 2016). It can for instance interfere with phenology, starch turnover and defence regulation, which may affect interactions with plant-associated organisms (Heinen 2021). As nights will most likely become significantly brighter, it is important to better understand the impacts on natural ecosystems (Rich & Longcore, 2013). Therefore, ALAN effects on plants and associated organisms warrant further studies (Heinen 2021).

Levels of ALAN will increase in the future because of the decreasing production costs, and the development of more energy- and cost-efficient light-emitting diode (LED) technology making light widely available and affordable (Hölker et al., 2010; Kyba et al. 2017). This will likely also increase the ecological consequences of ALAN. Although many forms of ALAN exist and are still in use, a clear trend towards LED technology has been observed in recent years. The most commonly used LEDs today are cool-white LEDs (with a colour temperature of >4200 K) and these are characterized by a narrow and high peak in the blue part of the irradiation spectrum, and a broad and lower peak in the red part of the spectrum. Cool-white LEDs can have devastating effects on many nocturnal animal species, and to mitigate these detrimental effects, several solutions have been suggested, including the use of different light colours (Van Geffen et al., 2014; Spoelstra et al., 2015), and to reduce light hours to periods that people are generally more active (International Darksky Association, www.darksky.org; Gaston et al. 2012; Day et al. 2015). A plausible scenario that could be implemented would be to strictly limit ALAN to the first part of the night, when human activity is still relatively high, and switching off the lights during the second half of the night. This would provide plants and animals with at least several hours of true darkness to recover. However, how ALAN and specific light schemes that reduce the number of light hours will affect plant defences and plant-herbivore interactions, is not known.

One key plant parameter that is affected by the quality, quantity and duration of light that a plant is exposed to is plant defence. First, a large body of knowledge describes how additional light, i.e., supplementing plants with artificial light *beyond* the daylight hours, influences plant defences. Generally, light supplementation into the dark hours increases plant defences. For example, tomato plants supplemented with low-density light at night have significantly higher immunity to pathogens compared to plants that grow in a dark night (Yang et al., 2015a). In another study, chickpea plants supplemented with light at night showed increased levels of the antioxidant ß-carotene (Wu et al., 2007), which may serve as a defence against oxidative stress (Mittler, 2002). One might conclude from this that a longer day leads to a more defended plant. A second body of scientific literature examines the effects of light reduction on plants, i.e., shading and light competition, on plant physiology, including defences, as partly mediated by changes in light intensity and spectral composition (e.g., Bennie et al. 2016; Smith et al. 2017; Grubisic et al. 2018). Light competition and shading can alter key processes in the jasmonic (JA) and salicylic acid (SA) signalling pathways (reviewed in Ballaré, 2009; Ballaré, 2014; Ballaré & Pierik, 2017), which is the backbone of plant defences against pathogens and insect herbivores. Reductions in light exposure, through competition and shading, result in decreased sensitivity to and suppressed JA and SA expression (de Wit et al., 2013; Ballaré & Pierik, 2017), although the SA-related processes are less well-understood (Ballaré & Pierik, 2017). Light competition and shade may also lead to reduced accumulation of plant secondary metabolites, and structural and indirect defences (Izaguirre et al., 2006, 2013; Engelen-Eigles et al., 2006; Moreno et al., 2009; Agrawal et al., 2013; Cargnel et al., 2014). Relevant to understanding the effects of ALAN on plants, one result of defence suppression under low-light or dark conditions is that plants are more susceptible to pathogens and herbivory at night (Roden & Ingle, 2009; Kraiselburd et al., 2017). This literature suggests that ALAN has the potential to affect plant defences but that this depends on the specifics of radiation at night.

ALAN may also affect interactions between plants and herbivores (Bennie et al., 2015; Yang et al., 2015b; Sanders et al., 2018; Heinen 2021). Different typical ALAN sources with different spectral characteristics have differential bottom-up effects on plant–aphid interactions (Bennie et al., 2015). For example, supplementing *Lotus pedunculatus* plants with amber ALAN, caused a decrease in densities of the aphid *Acyrthosiphon pisum* compared to no ALAN in the later stages of seasonal colony dynamics (Bennie et al., 2015). Importantly, even low light intensities at night can have significant effects, as ALAN can already suppress aphids under densities as low as 0.1 lux, which is far below commonly observed levels (Sanders et al., 2018). Although the mechanisms are not fully understood yet, it could be that ALAN results in systemic activation of defence mechanisms in the plant. For instance, in one study where tomatoes were supplemented with different spectral irradiance at night, particularly the plants exposed to red light (and to a lesser extent to other light colours) showed strong induced defence responses against the root-knot nematode *Meloidogyne incognita* (Yang et al., 2015b). These defence responses were accompanied by high levels of SA in root tissues, and upregulated transcript levels of SA-associated genes in the roots (Yang et al., 2015b). How irradiance levels associated with ALAN affects plant defences, however, is unknown.

In this study, we used a well-characterized plant–aphid model system, barley (*Hordeum vulgare* L.) and the English grain aphid (*Sitobion avenae* Fabricius), and exposed them to full nights of circa 20 lux ALAN (‘full-night’), half nights of circa 20 lux ALAN (‘part-night’) or no ALAN (‘control’) to investigate the effects of ALAN on plant–herbivore interactions and plant defence. We hypothesized that: (i) Exposure to ALAN reduces aphid colony growth compared to dark nights, mediated by an upregulation of defence responses. We predict that aphid colonization on plants will be negatively impacted by plants, through an upregulation of plant defences. (ii) Exposure to ALAN affects plant growth and health status, compared to dark nights. We predict that effects of ALAN will be positive, but likely minimal, as the levels of ALAN are expected to be too low to increase photosynthesis. (iii) A partial reduction in ALAN via a midnight elimination of light sources will reduce the hypothesized effects of ALAN, serving as a dark recovery time that benefits both plants and insects.

## Materials and Methods

### Soil

Soils for this experiment were derived from a former flower strip bordering an agricultural field of the Experimental Station Roggenstein (48.17989, 11.32030; altitude 524 m a.s.l.) at the Technical University of Munich. The soil was classified as sandy loam (50% sand, 33% silt, 14% clay, 2.4% SOM, pH 6.8, 1130 mg total-N/kg, 31.5 mg available-S/kg, 1.2 mg available-P/kg, 126 mg available-K/kg). The soil was sieved through a 2-cm mesh sieve to remove large stones, loam aggregates and roots.

### Plant–insect model system

Barley plants (*Hordeum vulgare*, cv. ‘Scarlett’) were used as model plants. This cultivar was selected as it was used in previous studies in our research group and because it was highly susceptible to aphids (Sanchez-Mahecha et al. 2022). As model aphids, we used the English grain aphid (*Sitobion avenae* ‘Fescue’, Hemiptera: Aphididae), a species that is specialized on grasses and is a common pest in cereal cropping systems. We have observed that this barley cultivar and aphid clone respond in terms of growth and population growth, respectively, to brief pulses of light disruption during the night in a pilot study. Aphid colonies have been maintained in a climate chamber at the institute at 15 °C (night) to 20 °C (day), under 16h:8h L:D conditions, on barley cultivar ‘Chanson’ for several years.

### Climate chamber conditions

This study was conducted in the TUM Model EcoSystem Analyzer (TUM*mesa*; Roy et al. 2021) in Freising, Germany. In this facility, eight separate high-performance climatic chambers allow for a very high level of between-chamber standardization of abiotic parameters and irradiation conditions (Teixeira et al., in prep.). Plants were grown under 16h:8h (L:D) daylength regimes, reflecting their summer growth period, with 21 °C and 16 °C temperatures during day and night, respectively. Daylights were turned on at 08:00, and were shut off at 00.00, which enabled us to sample under night conditions between 06:00 and 08:00. Relative air humidity was kept constant at 60%. During the day, plants received a full-spectrum light (Suppl. Fig. S1c), and the Photosynthetic Photon Flux Density (PPFD) was set to 500 μmol m^-2^s^-1^.

### Experimental design

We performed a full-factorial experiment involving ALAN treatment, two aphid treatments and four destructive (independent) harvesting days (Fig. 1). We used 16 biological replicates for ALAN control, 20 for part-night ALAN and 20 for full night ALAN treatment combinations, respectively (totalling 448 plants). In this study, we used seven climate chambers that were assigned a specific light treatment, and conducted the experiment in two separate runs, leading to 14 chambers (four assigned ALAN control, and five part-night and full-night ALAN, respectively). ALAN assignments to chambers were changed between runs (Supplementary Fig. S2) to control for climate chamber effects in our models. Biological replicates were divided over chambers and runs so that each chamber had four biological replicates of each combination (four replicates x eight combinations = 32 plants per chamber).

**Figure 1:**
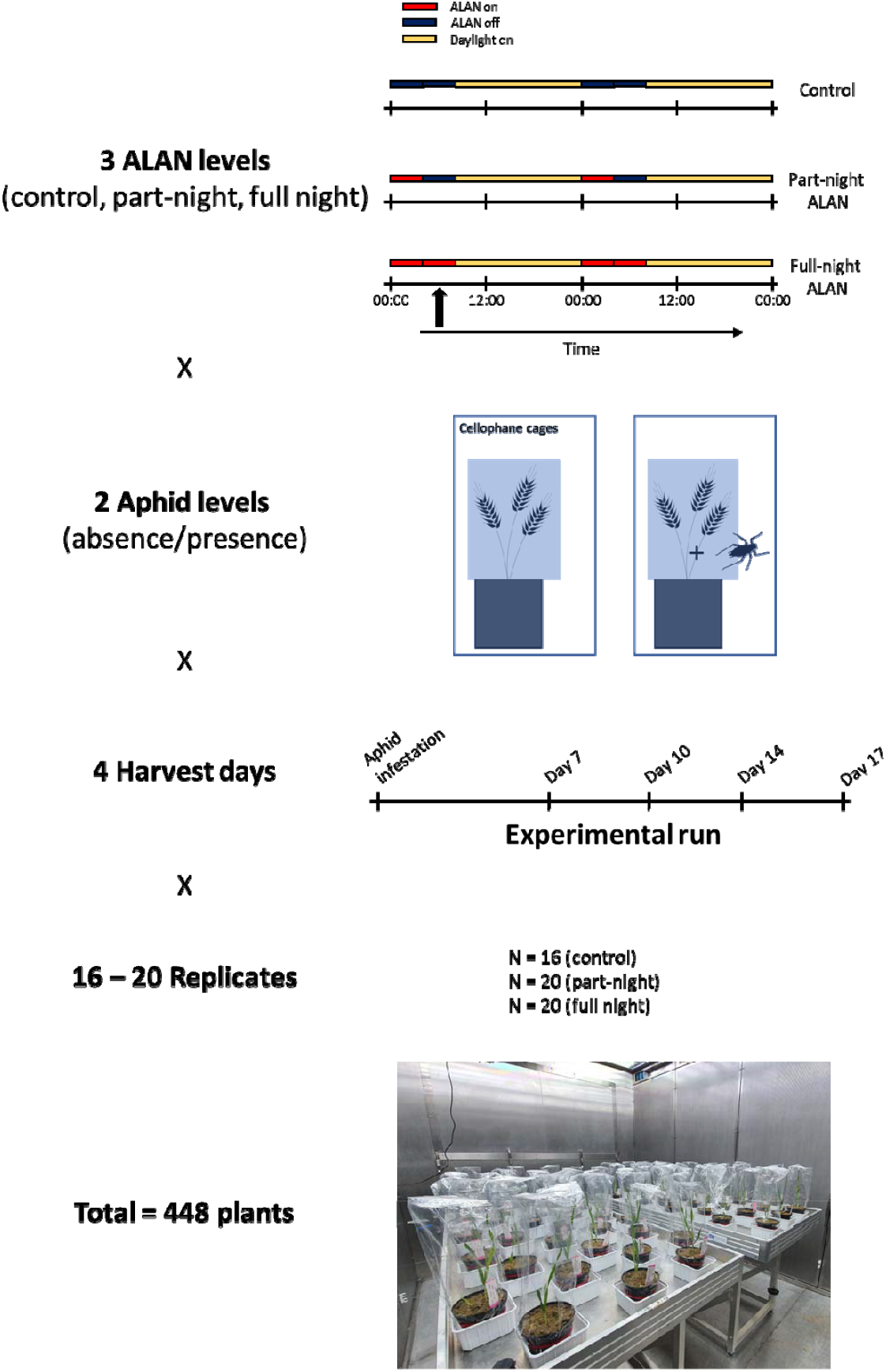
Schematic representation of the experimental design. The experiment was a full factorial combination of ALAN treatments, aphid treatment, temporal samplings, with replicates. ALAN treatment had three levels (control, part-night ALAN, full-night ALAN), two aphid levels (presence/absence), four destructive sampling dates, with 16–20 replicates each. In ALAN treatment scheme, red bars indicate ALAN lights on, dark blue bars indicate ALAN lights off, yellow bars indicate daylights on. The experiment was executed in seven climate chambers, over two experimental runs. Climate chambers were assigned specific ALAN treatments, and biological replicates were divided across assigned chambers and runs. ALAN controls had 16 biological replicates, divided over four different chambers, part-night ALAN and full night ALAN treatments had 20 biological replicates, divided over five different chambers. Each climate chamber contained 32 plants that were used for this study (pictured). The experiment had a total of 448 plants.

#### ALAN treatments

Each chamber was equipped with four linked LED modules each consisting of six white diodes per module (D-15-01-001, 2 W, 12 V, www.leds24.com), coupled to a dimmer to control illuminance. ALAN control chambers were kept dark at night, whereas part-night ALAN chambers were illuminated between 00:00 and 04:00. Full night ALAN chambers were lit with ALAN between 00:00 and 08:00. Illuminance was measured using a HR4000 spectrometer (using OceanView v1.6.7) paired to an Ocean Optics Custom Fiber Configuration OCF-102313 equipped with a cosine corrector with spectral diffusing material (Ocean Insight, Orlando, Florida, US) at pot level and illuminance was set to 15–20 lux, which are levels commonly observed for LED street lights (Bennie et al. 2016). Measurements with the same spectral meter underneath local LED street lights showed highly fluctuating and noisy signals of 5–100 lux. Under the experimental nocturnal light conditions, PPFD levels were below the measurable range of the fixed TUM*mesa* light sensors, indicating fewer than 0.5 μmol photons m^-2^ s^-1^. The spectral irradiance of the climate chambers during the day and spectral irradiance of the LEDs under maximum output capacity and under the dimmed ALAN treatment levels were measured and visualized after the experiment using an array-radiometer (Newport OSM400 200–800 nm, Sony CCD-Sensor 2046px and 50 μm gap, 1 nm spectral resolution, Newport Spectra Physics GmbH, Darmstadt, Germany). Supplementary Fig. S1 shows spectral irradiance for the LED modules under maximum capacity and under set ALAN treatments, and a visualization of a typical daylight spectral irradiance of the used climate chambers.

#### Plant and aphid treatments

Barley plants were grown with and without aphids in single round plastic pots (400 mL, 10 cm, Meyer, Göttingen, Germany). Pots were filled with soil, and watered with 50 mL of tap water and then left to acclimatize at room temperature for four days, then brought into the climate chambers, where they were left for another three days. Pots were checked daily and watered (20 mL) when the soil surface was dry. After this week of acclimatization, two surface-sterilized barley seeds (5% hypochlorite solution for 2 min, followed by a 1-min rinse with tap water) were planted in 1–2 cm deep holes made in the middle of the pot. Four days after sowing, most seedlings had germinated, and the smallest seedling was removed from each pot, so that only one plant per pot remained. Plants were checked daily, and watered to need, three times a week. One week after sowing, the height of each plant was recorded, and two pre-final (N4) instar individuals of *S. avenae* ‘Fescue’ were added to all pots assigned to the aphid treatment. The plants from both the aphid and no-aphid treatments were subsequently bagged around the pot rim with a permeable cellophane bag (Kopp Verpackungen, Höheischweiler, Germany), which was secured with elastic band to contain the aphids.

#### Temporal harvest

An *a priori* assigned subset of experimental plants from each combination was destructively sampled 7-, 10-, 14- and 17-days post-aphid infestation. This allowed us to follow aphid colony growth over time on independent plants. The destructive harvests were needed to sample for fresh plant material for gene expression.

#### Harvest and response variables

On the harvest days, total numbers of aphids were recorded. Among adult aphids, winged and unwinged aphids were distinguished. We measured plant height by measuring the distance between soil surface and the tip of the longest leaf. Chlorophyll levels were measured on the youngest fully expanded leaf at three points and recorded the average, using a chlorophyll meter (Konica Minolta SPAD-502Plus, Tokyo, Japan). Plants were placed on a cart in the evening, and the next morning, destructive sampling occurred between 06:00 and 07:00, before the daylights turned on. This ensured that we assessed the effects of the ALAN treatments – which by design occurred at night – on gene expression. Chambers were opened to remove plants for about 5–10 s. The harvest itself occurred in a dark corner of the corridor, with the only light coming from a dim emergency light above a door. Shoots were harvested within 5 min of removing plants from their chamber. To this end, the fresh shoots were clipped with sterilized scissors, folded in labelled aluminium foil, and flash-frozen in liquid nitrogen. This was done for the first three replicates for each treatment combination. We clipped and dried shoots of the remaining fourth replicate to determine shoot biomass. Roots for all four replicates per treatment combination were subsequently washed and dried to determine root biomass. Roots and shoots were oven-dried at 60 °C for at least three days before weighing.

#### Gene expression assay

In order to assess the effects of ALAN on plant-aphid interactions, we performed a reverse transcriptase-quantitative PCR on mRNA extracted from the frozen plant leaf material. For the gene expression assay we selected the plants harvested at day 10 post-infestation, as these plants did not yet show signs of aphid stress or yellowing, and hence provided high-quality samples for our gene expression assay. Before RNA extraction, three replicates from the same climate chamber were pooled for each treatment combination. Shoots of the three samples were ground and homogenized in liquid nitrogen. Effectively, the pooling resulted in 4–5 replicates for each ALAN-aphid combination (28 samples total). Total RNA was extracted from the homogenized sample using a QIAGEN RNeasy Plant Mini Kit (QIAGEN, Hilden, Germany), following the manufacturer’s protocol. RNA was converted to cDNA using SuperScriptII reverse transcriptase (Invitrogen, Thermo Fisher, Waltham, Massachusetts, US).

We selected four markers that are associated with defence-related proteins. Two markers amplify *PR1* and *PR17b*, two genes that code for pathogenesis-related proteins, and are generally involved in responses to biotrophic pathogens (Christensen et al. 2002; van Loon et al. 2006; Zhang et al. 2012), and are known to be upregulated in response to aphids (Grönberg 2006; Delp et al. 2009). Two other markers, *HvERF-like* and *HvWRKY22*, are associated with systemic defence against bacterial and fungal pathogens in barley (Dey et al. 2014; Lenk et al. 2019). The cDNA preparation was followed by qPCR on a 7500 Fast real-time qPCR system (Applied Biosystems, Thermo Fisher) with three technical replicates to obtain average Ct values per gene and sample. The qPCR was performed with the SensiMix SYBR low ROX kit (Bioline, Meridian Bioscience) using primers from Dey et al. (2014) and Shrestha et al. (2019).

For further statistical analysis, we determined delta Ct (δCt) values by *Ct_target_ - Ct_reference_*. Under the assumption that the endogenous reference gene has stable and high expression in all samples, indicated by a low Ct_reference_, this generally results in a positive δCt value for a sample, where a higher δCt indicates a lower expression. As we were interested in the relative expression of genes across treatments, the δCt values were used for further statistical analysis. However, since the experiment was replicated in two temporally separated runs, and this was an expected source of variation, we z-transformed the δCt values per run by subtracting the *run mean* δCt from each *sample* δCt, and divided this by the *standard deviation of the run mean* δCt, effectively taking out the run effect. The resulting z-score represents an effect size as the distance between run mean δCt and the sample δCt in units of standard deviation. As a positive δCt z-score indicates downregulation, which is visually not intuitive, we inverted the z-score signs, so that higher and lower relative gene expression match with the sign of the plotted values.

### Statistical analysis

All analyses were performed in R version 3.6.3 (R Development Core Team, 2020). Linear mixed models were performed using the package ‘lme4’ (Bates et al. 2015), and p-values were estimated using type II Wald Chi-Square tests using car::Anova from the ‘car’ package (Fox & Weissberg 2019). Data visualization was done using ‘ggplot2’ (Wickham 2016).

#### Plant variables

Plant variables were analysed using linear mixed models; as fixed factors we included ‘ALAN treatment’, ‘aphid treatment’ and ‘harvesting day’. We included ‘run’ as a covariate in the model, and ‘climate chamber’ as a random intercept, to correct for potential climate chamber effects.

#### Aphid variables

Aphid variables were analysed using generalized linear mixed models, with a Poisson error distribution; as fixed factors we included ‘ALAN treatment’ and ‘harvesting day’. We included ‘run’ as a covariate in the model, and ‘climate chamber’ as a random intercept, to correct for potential climate chamber effects. Only pots allocated to the aphid treatment were included in the analyses.

Significant effects of ALAN were further analysed using post-hoc Tukey tests using the ‘glht()’ command from the ‘multcomp’ package (Hothorn et al. 2008)

#### RT-qPCR

Gene expression data were analysed by means of z-transformed δCt values using linear mixed models. As fixed factors we included ‘ALAN treatment’ and ‘aphid treatment’, and ‘climate chamber’ as a random intercept, to correct for potential climate chamber effects.

We assessed relationships between gene expression and aphid numbers by averaging the aphid numbers on the samples that were pooled for the gene expression assays. The relationship between aphid numbers and inverted z-transformed δCt values was assessed by linear regression.

## Results

From the total of 448 sowed barley plants in the study, 430 survived until their assigned harvest and used for the analyses, of which 213 were assigned to aphid treatment, and 217 to no-aphid treatment. Across all treatment combinations 14-20 replicates were included in the plant-level analyses.

### ALAN and temporal effects on aphid colony growth

Aphid numbers increased with time, i.e., harvest day (Table 1, Fig. 2a). The total number of aphids per plant was significantly affected by ALAN (χ^2^_2_ = 102.5, p < 0.001, Fig. 2a). At the time of final harvest, aphid numbers on plants exposed to full night ALAN were 2.8% higher (mean 89.7 aphids) than those exposed to dark night controls (mean 87.4 aphids), whereas plants exposed to part-night ALAN had about 19.7% lower aphid numbers (mean 70.1 aphids) than controls. Part-night ALAN treatments had 21.9% lower aphid numbers than full-night ALAN treatments, and post-hoc Tukey tests indicated significant differences between all ALAN levels (Fig. S3a). We observed no significant interaction between harvest day and ALAN treatment in their effects on total aphid numbers (Table 1). When only adult aphids (winged and unwinged combined) were considered, numbers were affected by ALAN (χ^2^_2_ = 10.8, p = 0.005, Fig. 2b). At the time of harvest, adult aphid numbers were 9.8 and 32.2% lower on plants exposed to full night (mean 8.4 aphids) and part-night ALAN (mean 11.2 aphids) than on control plants exposed to dark nights (mean 12.4 aphids), respectively. Part-night ALAN treatments had 25.0 % lower adult aphid numbers than full night ALAN. Post-hoc Tukey tests indicated that at harvest time only part-night ALAN treatment was significantly lower from the two other levels (Fig. S3b). Harvest day also affected the number of adult aphids in the population (Table 1), as the number of adult aphids in the population increased over time (Fig. 2b). We observed no significant interaction between harvest day and ALAN treatment in their effects on adult aphid numbers (Table 1).

**Figure 2:**
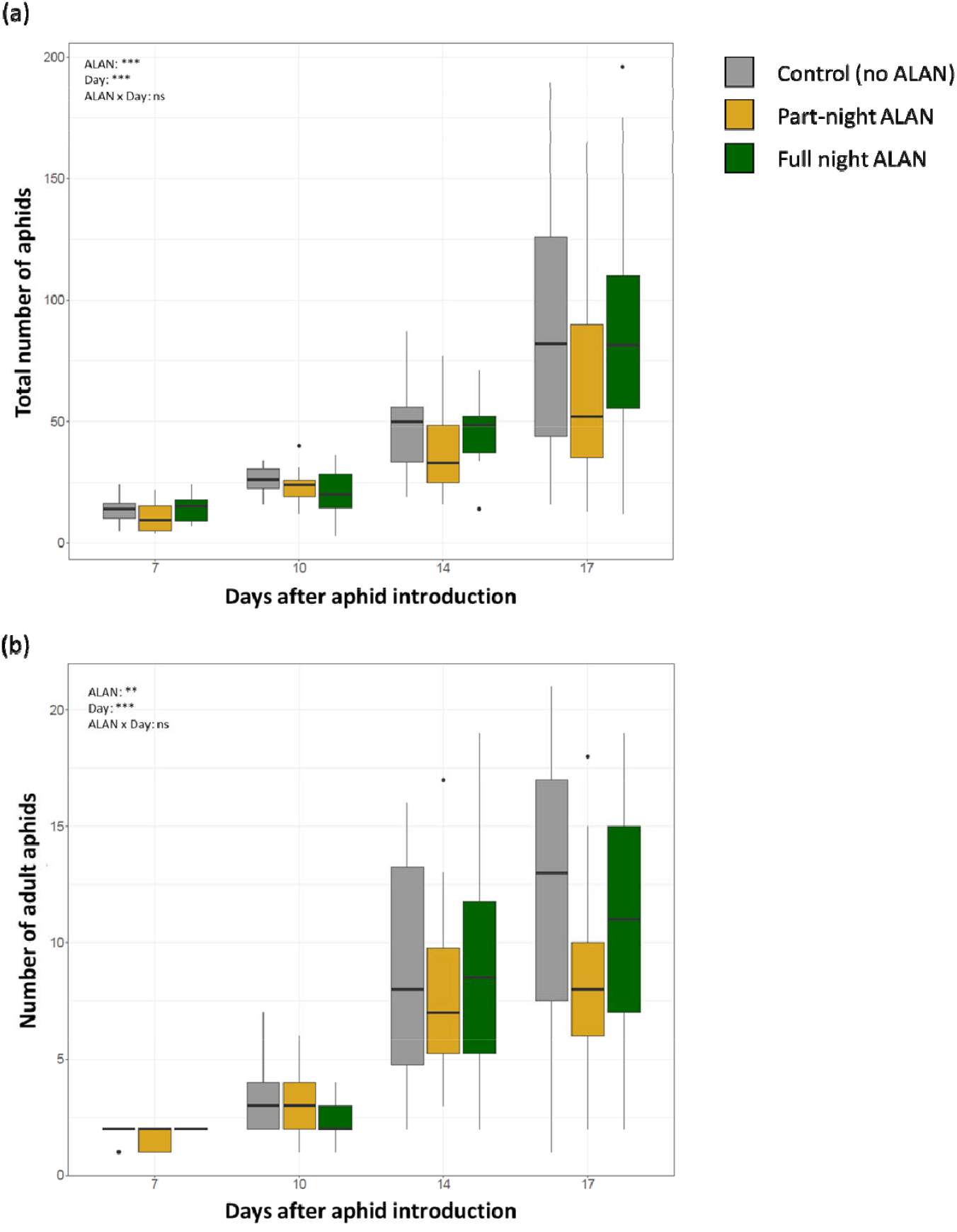
Box plots depicting effects of ALAN treatments and sampling time on a) aphid colony size (n = 16–20), and b) number of adult aphids within the population (n = 16–20). Box colours depict ALAN treatments, with grey being control, yellow part-night ALAN, and green full-night ALAN. Boxes represent median values with upper and lower quartiles, and whiskers represent 1.5x the interquartile range; individual dots represent outliers. Simplified mixed model output is indicated in the graphs with significance (ns = not significant, + p = 0.05 – 0.10, * p < 0.05, ** p < 0.01, *** p < 0.001) and detailed model output can be found in Table 1. Post-hoc Tukey tests were performed on a subset of the data on the final harvest point, and visualized separately for the subset in Fig. S3.

**Table 1:** Output of (generalized) linear mixed models testing the effects of Replicate Run, ALAN (A), herbivory (H) and time (T) on the number of total aphids, the number of adult aphids, plant height (cm), shoot biomass (g), root biomass (g), chlorophyll levels (SPAD units). Presented are degrees of freedom, Wald’s Chi-square statistics and p-values, obtained from the models using the car::Anova command in R. Significant effects (p < 0.05) are marked in bold, and marginally significant effects (0.05 < p < 0.1) in italics.

|  | Total aphids |  | Adult aphids |  | Plant height |  | Shoot dry mass |  | Root dry mass |  | Chlorophyll level |  |
| --- | --- | --- | --- | --- | --- | --- | --- | --- | --- | --- | --- | --- |
| | d.f. | $\chi^2$ (p) | d.f. | $\chi^2$ (p) | d.f. | $\chi^2$ (p) | d.f. | $\chi^2$ (p) | d.f. | $\chi^2$ (p) | d.f. | $\chi^2$ (p) |
| Run | 1 | **49.0 (<0.001)** | 1 | *3.1 (0.081)* | 1 | **64.4 (<0.001)** | 1 | **13.9 (<0.001)** | 1 | 0.7 (0.400) | 1 | **35.4 (<0.001)** |
| ALAN (A) | 2 | **102.5 (<0.001)** | 2 | **10.8 (0.005)** | 2 | *4.9 (0.085)* | 2 | *5.1 (0.076)* | 2 | **7.0 (0.030)** | 2 | **8.1 (0.017)** |
| Herbivory (H) | - | - | - | - | 1 | 0.6 (0.452) | 1 | 1.9 (0.166) | 1 | *3.7 (0.053)* | 1 | 2.7 (0.101) |
| Time (T) | 1 | **3073.6 (<0.001)** | 1 | **420.8 (<0.001)** | 1 | **119.3 (<0.001)** | 1 | **19.1 (<0.001)** | 1 | **4.4 (0.035)** | 1 | **64.7 (<0.001)** |
| A x H | - | - | - | - | 2 | 1.7 (0.438) | 2 | 0.1 (0.936) | 2 | 2.5 (0.284) | 2 | 1.5 (0.483) |
| A x T | 2 | 4.4 (0.111) | 2 | 3.2 (0.206) | 1 | 0.2 (0.923) | 1 | 2.7 (0.261) | 1 | 1.4 (0.486) | 1 | 1.3 (0.518) |
| H x T | - | - | - | - | 2 | 0.0 (0.984) | 2 | 2.7 (0.103) | 2 | 1.9 (0.167) | 2 | 0.2 (0.646) |
| A x H x T | - | - | - | - | 2 | 0.6 (0.738) | 2 | 0.7 (0.690) | 2 | 0.0 (0.992) | 2 | 1.3 (0.519) |

### ALAN, aphid and temporal effects on plant performance

Plant height and shoot dry biomass were only marginally (positively) affected by ALAN (plant height: χ^2^_2_ = 4.9, p = 0.084, shoot dry biomass: χ^2^_2_ = 5.1, p = 0.076) and plants tended to be higher under full night ALAN treatment (Fig. 3a,b), although post-hoc Tukey tests revealed no significant differences between the three ALAN levels across all experimental plants. Root dry biomass was significantly affected by ALAN (χ^2^_2_ = 7.0, p = 0.030, Fig. 3c), with post-hoc Tukey tests revealing that part-night ALAN significantly decreased root size compared to full night ALAN, but not to controls, across all experimental plants. Plant height and shoot biomass were not significantly affected by aphid treatment (Table 1), but aphids marginally reduced root biomass (χ^2^_1_ = 3.7, p = 0.053, Fig. 3c). Harvest day positively affected plant height, shoot and root dry biomass, as plants increased in size during the experiment (Table 1, Fig. 3). We observed no significant interactive effects between ALAN and herbivory on plant height, shoot or root biomass (Table 1). ALAN significantly affected chlorophyll levels in the plants (χ^2^_2_ = 8.1, p = 0.018), with plants exposed to full night ALAN exhibiting on average 9%, and plants exposed to part-night ALAN 4% higher chlorophyll levels than control plants exposed to dark nights, across all plants (Fig. 3b). Aphid presence did not significantly affect chlorophyll levels (Table 1), but these were significantly affected by harvest day, as levels decreased when plants grew older (Fig. 3b).

**Figure 3:**
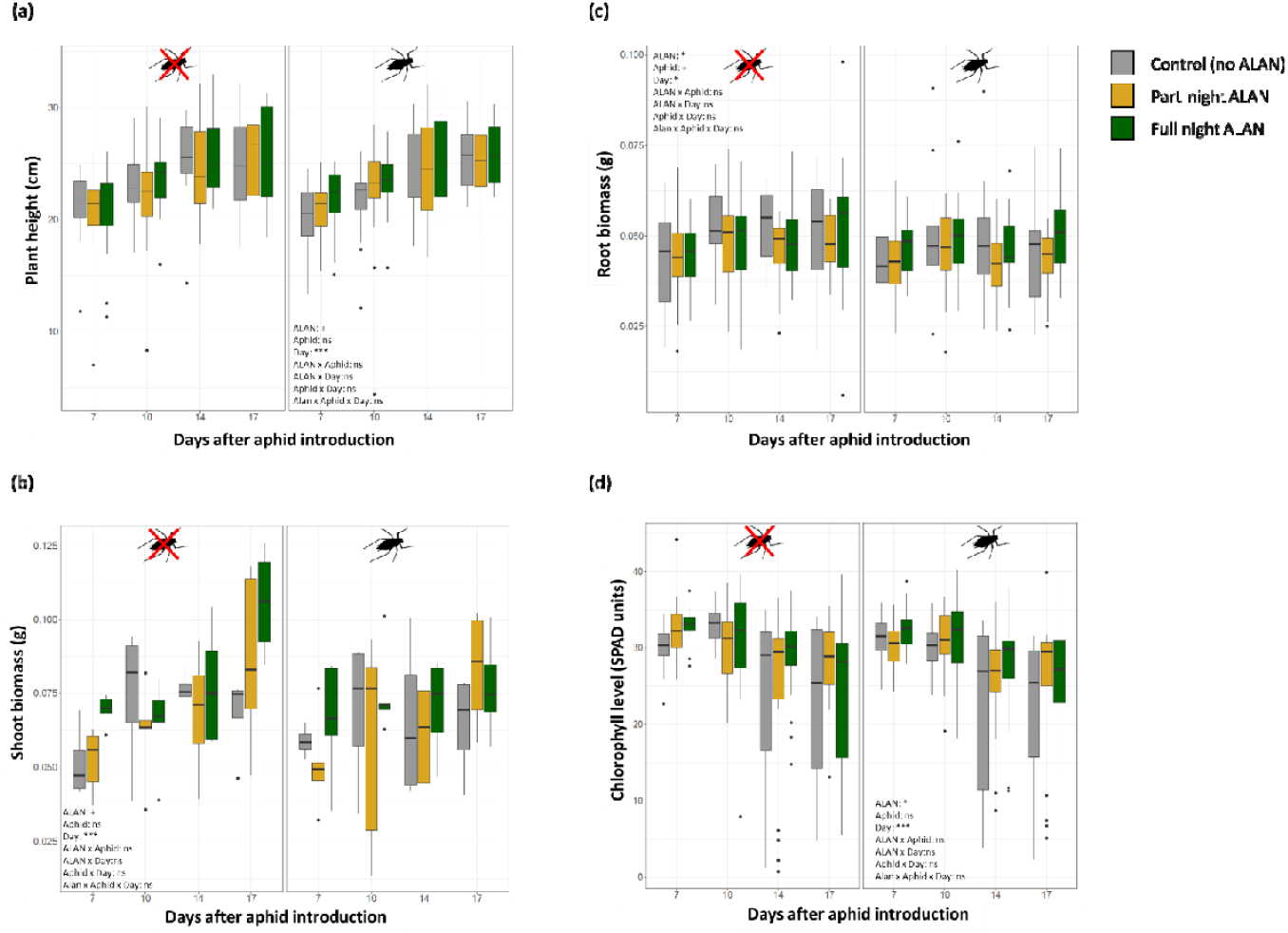
Box plots depicting effects of ALAN treatments and sampling time on a) plant height (n = 16–20), b) chlorophyll content (SPAD units) (n = 16–20), c) root (n = 16–20) and d) shoot dry biomass (n = 4–5). Box colours depict ALAN treatments, with grey being control, yellow part-night ALAN, and green full-night ALAN. Boxes represent median values with upper and lower quartiles, and whiskers 1.5x the interquartile range; individual dots are outliers. Simplified mixed model output is indicated in the graphs with significance (ns = not significant, + p = 0.05 – 0.10, * p < 0.05, ** p < 0.01, *** p < 0.001), and detailed model output can be found in Table 1.

### ALAN and aphid effects on plant gene expression

The expression of some defence genes was marginally or significantly affected by the ALAN treatments while the presence of aphids marginally or significantly increased expression of most defence genes (Table 2). The expression of *HvWRKY22* was marginally affected by ALAN treatments (χ^2^_2_ = 5.9, p = 0.053, Fig. 4a), with a trend for expression to be downregulated when plants were exposed to full night or part-night ALAN treatments (Fig. 4a). The expression of *HvWRKY22* was strongly upregulated in aphid treatments (χ^2^_1_ = 8.0, p = 0.005, Fig. 4a). No interactive effects were observed on the expression of *HvWRKY22* (Table 2). The expression of *HvPR1* was marginally upregulated by aphid treatments (χ^2^_1_ = 3.6, p = 0.057, Fig. 4b), but effects of ALAN treatments or interactive effects on the expression of *HvPR1* were not observed (Table 2). The expression of *HvPR17b* differed significantly between ALAN treatments (χ^2^_2_ = 10.1, p = 0.006, Fig. 4c) and was marginally upregulated by aphid treatments (χ^2^_1_ = 3.1, p = 0.077, Fig. 4c). Post-hoc Tukey tests indicated that expression of *HvPR17b* was significantly lower when plants were exposed to full night or part-night ALAN, compared to ALAN controls across all plants. The expression of the *HvERF-like* gene was not significantly affected by any of the treatments (Fig. 4d).

**Figure 4:**
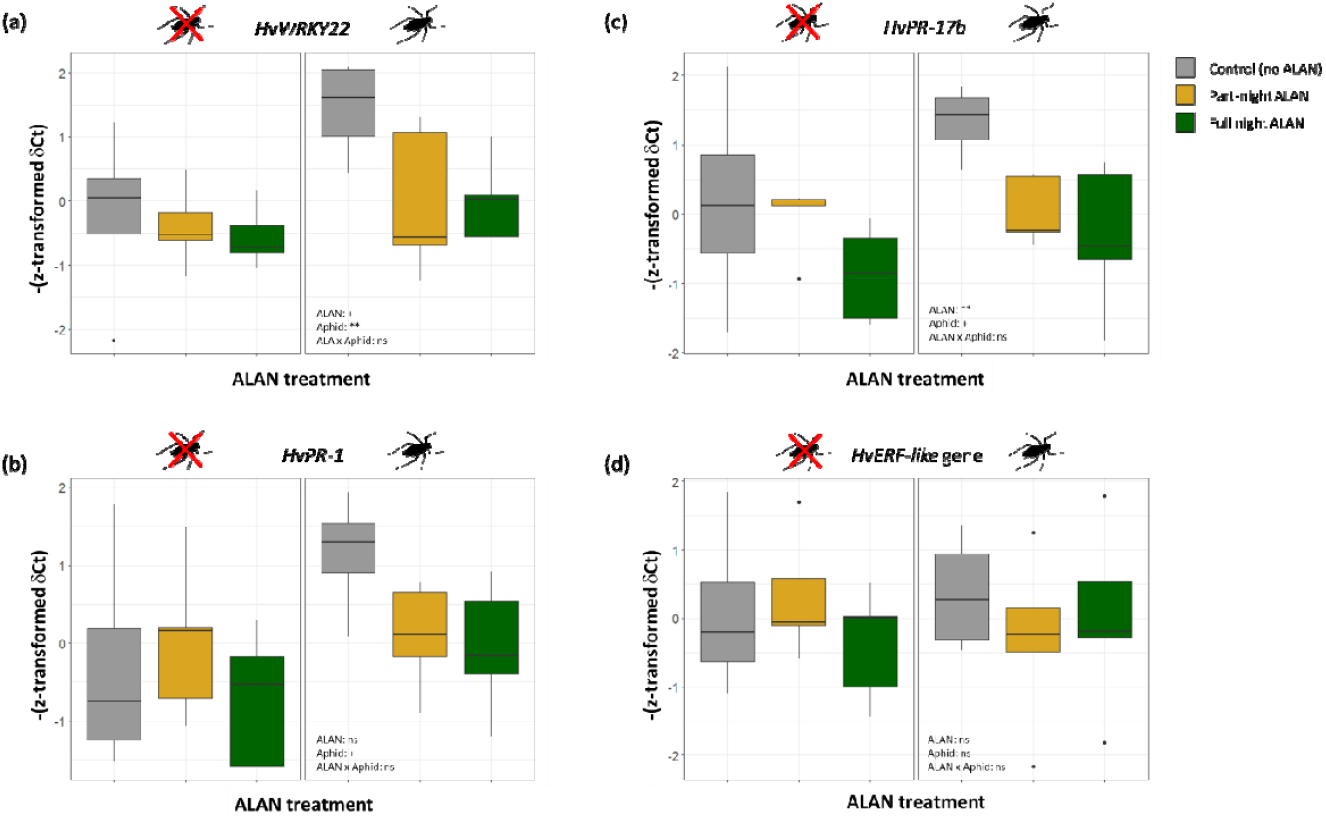
Box plots depicting effects of ALAN treatments on standardized gene expression of a) *HvWRKY22*, b) *HvPR-1*, c) *HvPR-17b* and d) *HvERF-like gene* markers (each n = 4 (control) or 5 (treatment)). Box colours depict ALAN treatments, with grey being control, yellow part-night ALAN, and green full-night ALAN. Boxes represent median values with upper and lower quartiles, and whiskers represent 1.5x the interquartile range; individual dots represent outliers. Simplified mixed model output is indicated in the graphs with significance (ns = not significant, + p = 0.05 – 0.10, * p < 0.05, ** p < 0.01, *** p < 0.001), and detailed model output can be found in Table 2.

**Table 2:** Output of linear mixed models testing the effects of ALAN (A), herbivory (H) and time (T) on the standardized expressi on of *HvWRKY22, PR-1, PR-17b*, and *HvERF-like* marker genes. Expression data was standardized by replicate run, which was therefore excluded from the model. Presented are degrees of freedom, Wald’s Chi-square statistics and p-values, obtained from the models using the car::Anova command in R. Significant effects (p < 0.05) are marked in bold, and marginally significant effects (0.05 < p < 0.1) in italics.

|  | *HvWRKY22* |  | *PR-1* |  | *PR-17b* |  | *HvERF-like* |  |
| --- | --- | --- | --- | --- | --- | --- | --- | --- |
| | d.f. | $\chi^2$ (p) | d.f. | $\chi^2$ (p) | d.f. | $\chi^2$ (p) | d.f. | $\chi^2$ (p) |
| ALAN (A) | 2 | *5.9 (0.053)* | 2 | 3.3 (0.192) | 2 | **10.1 (0.006)** | 2 | 0.5 (0.779) |
| Herbivory (H) | 1 | **8.0 (0.005)** | 1 | *3.6 (0.057)* | 1 | *3.1 (0.077)* | 1 | 0.0 (0.998) |
| A x H | 2 | 3.5 (0.170) | 2 | 2.4 (0.302) | 2 | 1.9 (0.381) | 2 | 1.3 (0.531) |

The total number of aphids on those plants used for qPCR analyses positively correlated with the expression of *HvWRKY22* (R^2^ = 0.31, F_1,12_= 5.5, p = 0.037; Fig. 5a), *HvPR1* (R^2^ = 0.55, F_1,12_ = 14.8, p = 0.002; Fig. 5b), and *HvPR17b* (R^2^ =0.71; F_1,12_ = 29.4, p < 0.001,; Fig. 5c), but not with the *HvERF-like* gene (R^2^ = 0.002, F_1,12_ = 0.02, p = 0.891; Fig. 5d).

**Figure 5:**
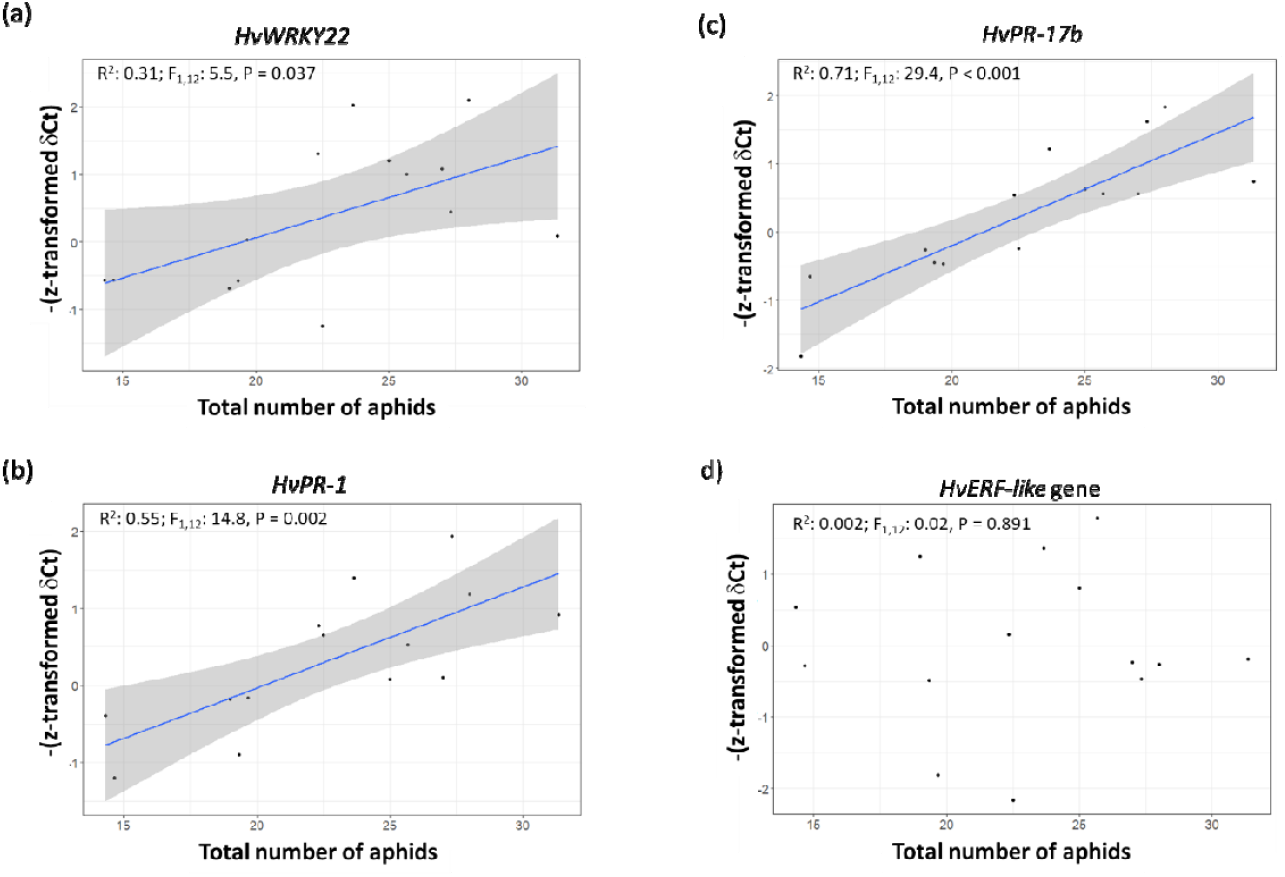
Relationships between the average number of aphids on barley plants assigned to a pooled sample, and the standardized gene expression of a) *HvWRKY22*, b) *HvPR-1*, c) *HvPR-17b* and d) *HvERF-like gene* markers across ALAN treatments. Linear correlations with 95% confidence intervals were fit if the linear relationship was significant.

## Discussion

In this study, we exposed plants and associated aphids to either full nights of ALAN, nights with a part-night ALAN treatment, where ALAN was switched off halfway through the night, or controls with completely dark nights. We found that plant growth was only minimally affected by ALAN, whereas aphid colony growth and the number of adult aphids in the colony were impacted significantly by ALAN treatments. Importantly, we found strong differences between full and part-night ALAN treatments. Contrary to our hypotheses, ALAN mitigation – by limiting the duration of light exposure at night – had stronger negative effects on aphid colony growth than nights with full exposure to ALAN. Gene expression assays revealed that defence-associated gene expression was affected by ALAN and aphid presence in three of four genes, but in these three genes, expression positively correlated with aphid colony size, suggesting that the effects of ALAN on plant defences are likely regulated through indirect effects of ALAN on plants via direct disruptive effects on aphid colonies, rather than via regulation by the plant in direct response to ALAN.

In agreement with our first hypothesis, exposure to ALAN suppressed aphid colony growth, although particularly in the full-night ALAN treatment, we observed a recovery of aphid numbers over time. Previous studies show suppressive ALAN effects on plant–aphid interactions in other model systems, where aphid suppression was associated with changes in plant phenology, and increases in aphid parasitism by parasitoids (Bennie et al. 2015; Sanders et al. 2015; Bennie et al. 2018; Sanders et al. 2018). Importantly, our study shows that aphid colony decline also takes place without such external confounding factors. Contrary to the present study, a previous field experiment with *H. vulgare* and *S. avenae* detected no effects of ALAN on final aphid densities at nine weeks (Sanders et al. 2018), indicating that the strongest effects may be early after colonization, and may disappear over time, which is in line with our observations that aphid numbers recover in full night ALAN treatments. However, it should be noted that the referenced study included parasitoids, which as aphid antagonists may have affected aphid colony sizes. Other studies that have assessed ALAN effects on aphid colony size over time have generally included repeated measures of the same colony (Bennie et al. 2015; Sanders et al. 2015), which allows to assess individual colony responses, but also elevates the risk of including colony-specific effects that persist over time. In our study, aphid colony growth was assessed over time on independent samples which provides a higher temporal resolution, and also reinforces the generality of ALAN impacts on aphid colony formation, especially during early colonization.

An important question is what drives the ALAN-induced impacts on aphid numbers? – Two answers are plausible: On the one hand, ALAN could influence aphids indirectly, via changes in plant nutritional quality or quantity (plant defence compound and nutritional quality or plant size), which commonly affects insect performance through bottom-up effects (Will & Van Bell 2006). On the other hand, ALAN may directly impact aphids, through disturbance of natural behaviour such as settling times that may limit feeding, and ultimately, fitness. Our data suggest that plant size was only marginally affected by ALAN. Although various studies have suggested that resource availability could be the main driver through which ALAN impacts insect responses (Sanders et al. 2015; Bennie et al. 2015), plant size under levels of infestation observed in our study are unlikely to be limiting to aphid population growth, as plants were large enough to sustain the relatively small aphid numbers. It thus seems more plausible that potential bottom-up effects are mediated via plant quality, not quantity. We observed a significant increase in chlorophyll levels – a proxy for plant health and quality – when plants were exposed to ALAN, but as our data suggested only a weak and positive relationship with aphid numbers, it is highly unlikely that chlorophyll levels alone are responsible for aphid suppression under ALAN. Future research should focus on how ALAN may affect plant quality for aphids, and especially the diurnal variation in the phloem composition (Douglas 2006), in order to better understand the possible links between ALAN, the plant and ALAN-induced aphid suppression.

We assessed the expression of four defence-associated genes in barley corresponding with different defence pathways, to test whether exposure of plants to ALAN would result in a regulation of specific pathways. We measured expression of two *PR* genes, which are commonly associated with innate immunity in plants (van Loon et al. 2006; Wang et al. 2018). In particular, *HvPR1* and *HvPR17b* are induced upon infection of barley with pathogens and are presumed to positively correlate with defence (Christensen et al. 2002; Grönberg 2006; Delp et al. 2009; Zhang et al. 2012). Further, *HvWRKY22* and *HvERF-like* have previously been positively associated with SAR-like systemic defence in barley (Dey et al. 2014). We observed a marginally significant effect (p = 0.053) of ALAN on the expression of *HvWRKY22*, and a strongly significant effect of ALAN on the expression of *HvPR17b*. We did not find any effect of ALAN or aphid treatments on the expression of *HvERF-like* gene expression, suggesting that this gene, which is associated with systemic pathways (Dey et al. 2014), is unlikely to respond to variation in light at night and is not involved in plant defences against aphids.

Contrary to our expectation that ALAN would upregulate plant defences, ALAN decreased the expression of *HvWRKY22* and *HvPR17b*, and we observed similar but non-significant patterns in *HvPR1*. In fact, for *HvWRKY22*, *HvPR1* and *HvPR17b*, but not for *HvERF-like* gene, the gene expression correlated tightly and positively with the aphid numbers present on the plant. This suggests that patterns of expression differing between ALAN treatments were likely not the result of direct ALAN effects on plant defence regulation, but instead more likely a result of direct disruptive effects of ALAN on aphid feeding or settling behaviour and resultant suppression of colony growth, and that this, in turn, is reflected by a reduced transcript accumulation of certain defence-associated genes. Importantly, from our results, we cannot conclusively state that ALAN does not affect plant defences. Our analyses focused on a limited number of genes that are involved in constitutive and induced plant defence responses against aphids. A more holistic approach, for instance using meta-transcriptomics of plants under ALAN or dark night treatments could reveal how ALAN may affect transcriptional regulation of phytohormonal defence mediation, as well as other pathways involved, for instance, in secondary metabolism.

There are various alternative plant physiological pathways through which ALAN might affect the phytobiome (Heinen 2021). For instance, exposure to ALAN can disrupt stomatal activity, above-belowground allocation, and the nocturnal starch turnover from above-to below-ground compartments (Stitt & Zeeman, 2012; Kwak et al. 2017; 2018). Disruptions in starch metabolism may result in changes in concentration and composition of sugars in phloem tissues, and as aphids are directly affected by phloem sugar concentrations via the ‘sugar barrier’, which requires strong osmoregulation in the aphid gut to excrete it (Douglas 2006), it is entirely plausible that such a pathway might affect probing, feeding and reproductive success, leading to altered aphid colony growth. As such, future studies should investigate ALAN effects on sugars, amino acids, micronutrients and secondary metabolites in the phloem that are important for aphid performance. For instance, changes in carbon metabolism over diurnal cycles can greatly impact interactions between plants and their microbiome (Baraniya et al. 2018), and knowing how ALAN may impact on carbon metabolism and affect interactions between plants and other organisms will be an exciting direction for future research (Heinen 2021).

ALAN can also directly impact on aphid colony growth, via effects of light on aphid behaviour. Aphids perceive light intensity and can respond to low light intensities by producing winged adults (Alkhedir, Karlovsky & Vidal 2010). Aphids also express diurnal rhythmicity in locomotor and metabolic activity (Beer et al. 2017), even independent of their plant hosts, which are strongly rhythmic (Joschinski et al. 2016). Despite strong cyclic circadian clock gene expression, aphids continue the various stages of their feeding cycles throughout the night, which is hypothesized to be an important element for osmoregulation of high phloem sugar content (Douglas 2006; Nalam et al. 2021). Light has been shown to disrupt aphid behaviour, and more strongly so during the night (Narayandas & Alyokhin 2006). This effect is likely stronger in plants with more open canopies, where light reaches the insect directly. We speculate that ALAN led to disruptions of settling and feeding behaviour, lowering the uptake of resources, which in turn explain smaller aphid colony sizes and number of matured adults, and the tightly linked defence gene expression, as observed under our treatments. Further tests to examine how ALAN impacts aphid feeding behaviour, should make use of electrical penetration graph analyses (Tjallingii 1978), which record the probing behaviour of aphids over time, to compare aphid behaviour under dark and lit nights.

An important finding is that contrary to our hypothesis, a mitigation of ALAN levels at night through a midnight shutoff of the light source enhanced adverse ecological effects of ALAN on insects. A popular belief is that the ecological side effects of light pollution could be mitigated by optimizing lighting schemes to match human activity patterns, and turning off lights after peak activity. However, our data suggest that after experiencing a short duration of ALAN, turning off the lights has more detrimental effects than keeping them on. These findings are difficult to explain. One possibility is that a midnight shutoff strategy may be experienced as a double ecological stressor. First, the organisms experience a lit night, which may be seen as the first stressor, which is then followed by a severely reduced dark recovery period, i.e., further stress, although the mechanism is unclear. However, we argue that the ecological consequences of specific mitigation strategies need to be studied before they are readily implemented, as despite good intentions, they may sometimes have unexpected side effects (Cieraad et al. 2022).

Our study also highlights the physical limitations of ALAN research. One example is the typical choice to use illuminance – expressed in lux, which is a photometric unit – in most ALAN research. This means, that the radiometric quantity will be weighted with a visual function describing the wavelength dependent response of the human eye. The photopic function of the human eye have the maximum at 550 nm, while 380–780 nm is named as ‘light’ (Aphalo et al. 2012). However, light is in the physical sense better described as radiant energy per time per effective receiver surface and wavelength interval. This is where ALAN research becomes more problematic, as its spectral irradiance generally falls below thresholds measurable by standard equipment. Another problematic aspect is that in the lower ranges of spectral irradiance, the ratio of noise to signal is very high (cf. Suppl. Fig. S1b). Measurements conducted under LED street lights in the field confirm low signals of irradiance, with comparatively high levels of noise. A better physical characterization of ALAN irradiance spectra, and its reach in nature is essential to understand the breadth of ecological impacts, especially on sessile organisms such as plants.

Despite significant advances in the field, the effects of ALAN on plants and plant-associated organisms are understudied (Heinen 2021). We aimed to generate a better understanding of how ALAN impacts on aphid colony growth over time, as mediated via plant quality and quantity. In general, ALAN impacts on barley plants appear to be minor, but regardless exposure to ALAN leads to substantial effects on aphid colony growth, which appear to be direct disruptive effects of ALAN, rather than plant-mediated effects. Our study especially highlights the difficulties associated with mitigation of the ecological side effects of ALAN, as a reduction of the light intervals increased the negative impact of ALAN on aphids. Rather little is known about how ALAN affects insect communities on crops and wild plant species. These findings highlight the need to study the impacts of ALAN in natural ecosystems, and we show that the solution to light pollution may be more complex than the flip of a switch.

## Supporting information

Supplementary figures to Heinen et al.

## Acknowledgements

We thank Roman Meier for advice and engineering support in the TUM*mesa* phytochamber facilities. We thank Hans Hausladen and Stefan Kimmelmann for advice and help in obtaining soil for the pot experiment. We thank Sharon Zytynska for supporting the project. The experiment was supported by DFG grant 397565003 WE 3081/36-1 awarded to Sharon Zytynska and WWW.

