## Supplementary figures to Heinen et al. for "Part-night exposure to artificial light at night has more detrimental effects on aphid colonies than fully lit nights"

Supplementary Fig. S1: Spectral irradiance graphs of a) the experimental LED modules at maximum capacity, measured 5 cm from the source, b) the experimental LED modules dimmed to experimental ranges of 15–20 lux, measured at pot level, and c) one of the climate chambers used in the experimental setup. Note that spectral irradiance was measured after the experiment was terminated, and at this time the climate chamber was set to slightly higher irradiance levels, than was the case during the experiment. Although spectral irradiance in the experimental phase were lower, the realized spectra were highly similar.

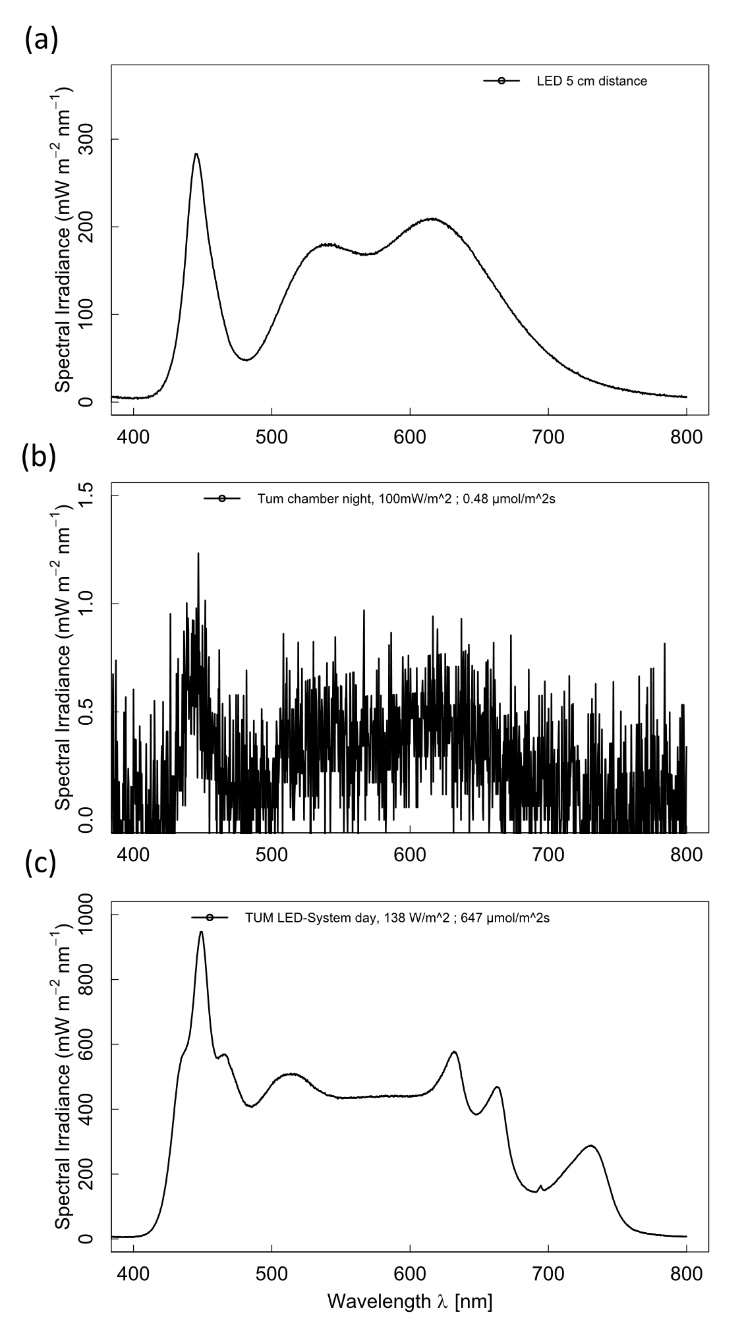

Supplementary Fig. S2: Overview of the technical replicates of ALAN treatment to climate chambers to ALAN treatments in the two experimental runs. In addition to the switching of treatments assigned to chambers between runs, our statistical models contained chamber identity as a random effect.

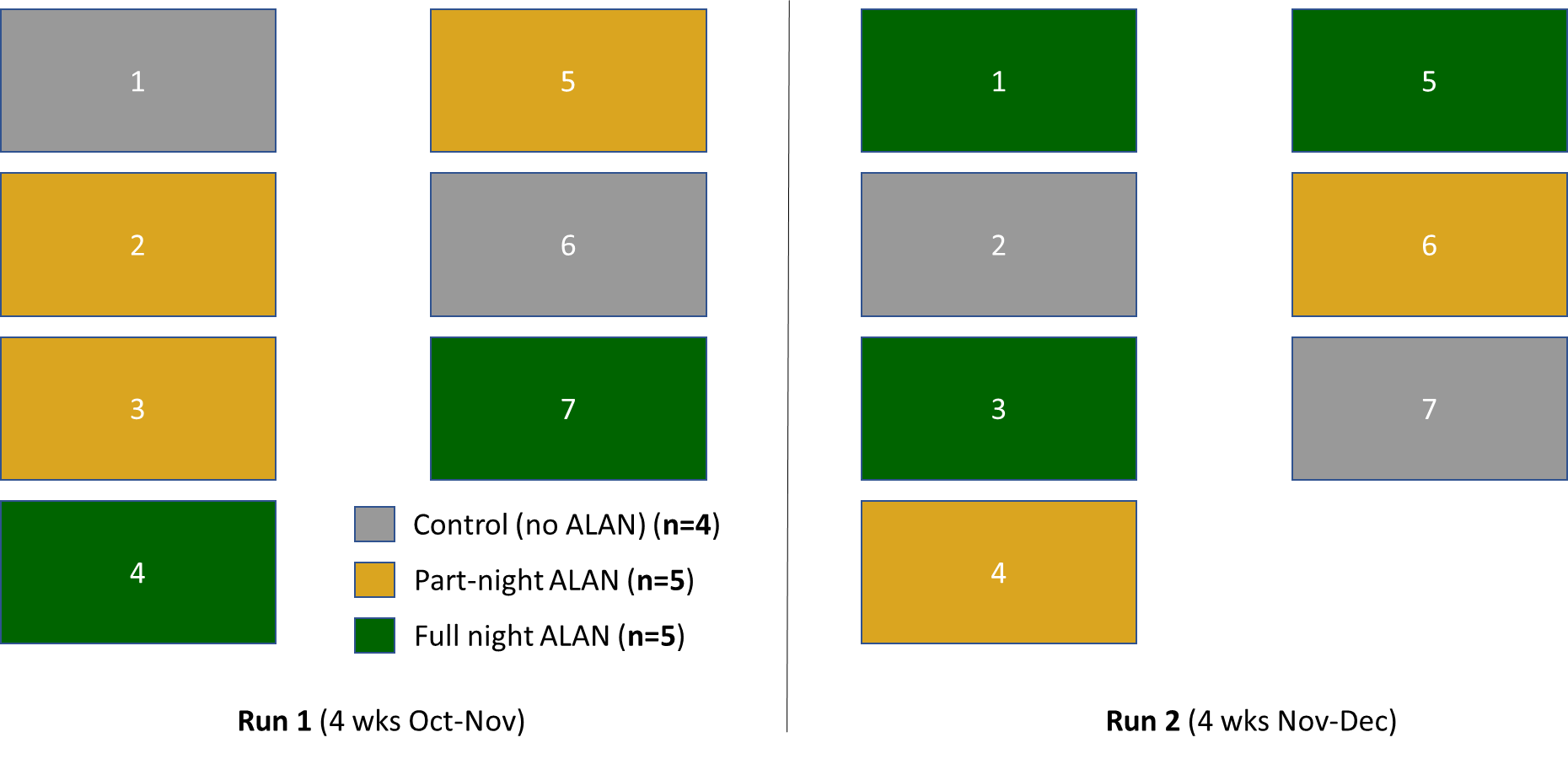

Supplementary Fig. S3: Box plots depicting effects of ALAN treatments on a) aphid colony size (n=16-20), and b) number of adult aphids within the population (n=16-20), at the time of harvest, 17 days post-infestation. Box colours depict ALAN treatments, with grey being control, yellow part-night ALAN, and green full-night ALAN. Boxes represent median values with upper and lower quartiles, and whiskers represent 1.5x the interquartile range; individual dots represent outliers. Means are symbolized by triangle shapes inside the boxes. Post-hoc Tukey tests were performed using the glht() command in the ‘multcomp’ package, and different letters indicate significantly different means.

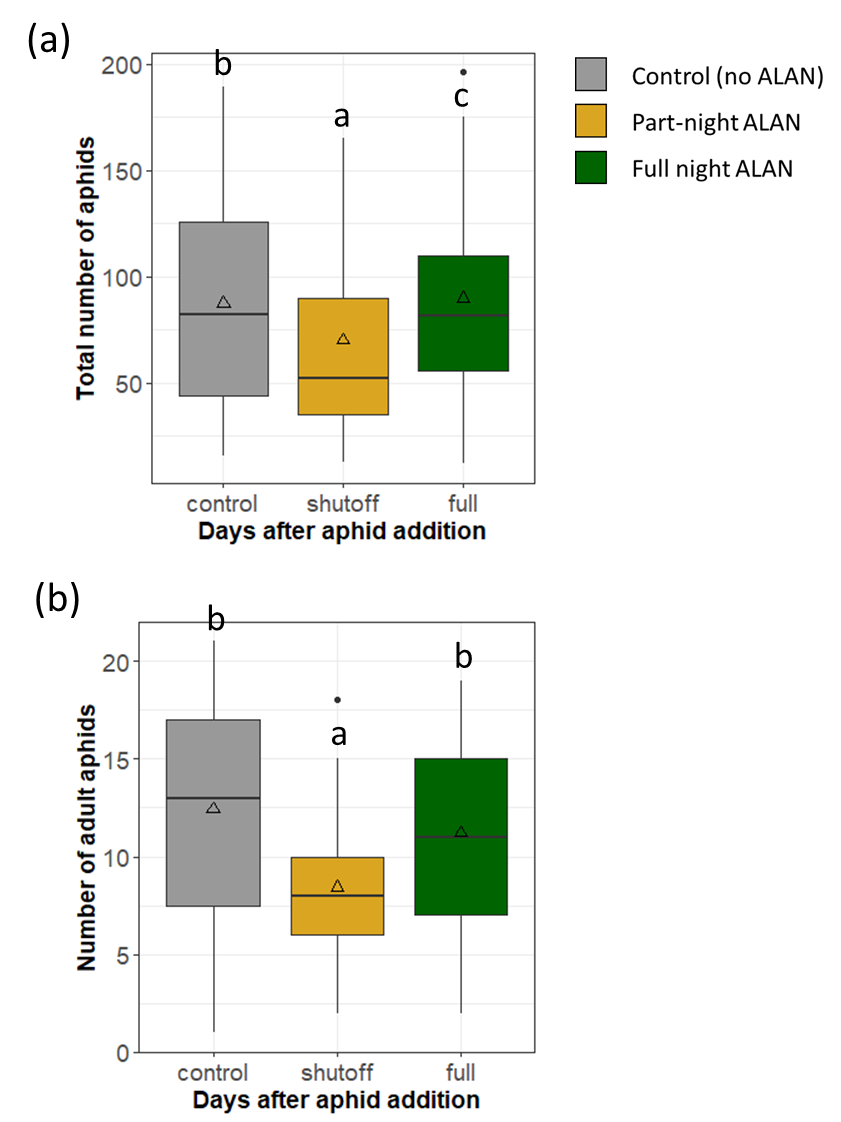
